# Drug combination prediction for cancer treatment using disease-specific drug response profiles and single-cell transcriptional signatures

**DOI:** 10.1101/2022.03.31.486602

**Authors:** Daniel Osorio, Parastoo Shahrouzi, Xavier Tekpli, Vessela N. Kristensen, Marieke L. Kuijjer

**Author notes:** These authors contributed equally to this work (Daniel Osorio, Parastoo Shahrouzi). Correspondence &.

## Abstract

Developing novel cancer treatments is a challenging task that can benefit from computational techniques matching transcriptional signatures to large-scale drug response data. Here, we present ‘*retriever*,’ a tool that extracts robust disease-specific transcriptional drug response profiles based on cellular response profiles to hundreds of compounds from the LINCS-L1000 project. We used *retriever* to extract transcriptional drug response signatures of triple-negative breast cancer (TNBC) cell lines and combined these with a single-cell RNA-seq breast cancer atlas to predict drug combinations that antagonize TNBC-specific disease signatures. After systematically testing 151 drug response profiles and 11,325 drug combinations, we identified the combination of kinase inhibitors QL-XII-47 and GSK-690693 as the topmost promising candidate for TNBC treatment. Our new computational approach allows the identification of drugs and drug combinations targeting specific tumor cell types and subpopulations in individual patients. It is, therefore, highly suitable for the development of new personalized cancer treatment strategies.

## Introduction

Developing new drugs and pharmacological regimens to treat complex diseases such as cancer is an expensive and challenging task (1). Several computational techniques that use structural interactions (2), transcriptional signatures (3–5), biological networks perturbations (6, 7), and data mining (8) are available to aid in the discovery of new treatment regimens which could improve patient management (9). Methods based on transcriptional signatures use the observed changes in gene expression profiles between samples from patients affected with the disease under study compared to samples from control subjects, and match these to the response profiles of cell lines that act as surrogates for the disease to different compounds. These data can be combined to identify compounds that may antagonize the disease’s transcriptional changes, returning it to a more healthy-like state (5). Finding specific transcriptional response profiles to drug candidates, as well as robust disease-associated transcriptional alterations, is therefore crucial for such approaches to produce reliable predictions.

To obtain precise disease-specific transcriptional signatures that can be targeted by pharmacological compounds, one needs to account for transcriptional variability between patients by including samples from multiple individuals (10). This allows for the identification of a target set of genes that is constitutively expressed with the same magnitude and pattern across patients/cells. Targeting such a consistently expressed set of genes is thought to increase the effectiveness and safety of the treatment (11). Transcriptional signatures are measurable at a variety of resolutions, ranging from low resolution, as in the complex mixture of cells present within tissues, to high resolution, such as single-cells (12). The Cancer Genome Atlas (TCGA) project provides RNA-sequencing (RNA-seq) data for tumors and adjacent tissues (13). While such tissue-level data have been useful in the identification of cancer subtypes and prognostic signatures, the identification of accurate transcriptional signatures between disease and healthy states has been hindered by the wide heterogeneity of cell types found in tumor tissues (14).

This issue can be overcome by using single-cell RNA-seq data (15). A drawback of single-cell RNA-seq experiments is that, due to the higher cost of profiling of single cells, often fewer biological replicates are available. However, by combining multiple experiments, sample sizes of cells with a desired molecular phenotype can be increased, improving characterization of the transcriptional changes observed in disease (16). In addition, the construction of pseudo-bulk profiles (the sum of the expression values of a gene across all cells obtained from the same individual) from single-cell data makes it possible to account for gene expression variability among patients (17, 18).

The LINCS-L1000 project measures 978 “landmark” transcripts with a low-cost bead-based assay and computationally infers the expression of 11,350 additional genes (4, 19). Its Phase I release (GEO GSE92742) contains 473,647 signatures of 20,413 small molecules, 18,493 shRNAs, 3,492 open reading frames and 622 ligands across 76 cell lines, although coverage is uneven: only 2,834 compounds were profiled in ten or more cell lines. These profiles have previously been used to identify potential drugs that can be repurposed to treat a variety of diseases, including cancer (3). The compounds were not selected in relation to any disease; query tools such as the CLUE platform rank perturbagens by a connectivity score that measures how similar or opposite their signatures are to a user-supplied set of up- and down-regulated genes (4, 20). However, these predictions are not based on robust tissue- or disease-specific transcriptional profiles and may therefore over- or underestimate the potential effect of drugs on specific diseases. Additionally, the LINCS-L1000 web portal returns an exhaustive list of independent experiments with matching inverse patterns, rendering it difficult to identify compounds that induce stable transcriptional responses in cell lines derived from the same disease across multiple time points and drug concentrations. Thus, being able to extract robust disease-specific transcriptional drug response signatures that are consistent at different time points, drug concentration, or cell line from the profiles provided by the LINCS-L1000 project would significantly improve drug prioritization and accelerate the identification of new pharmacological options for personalized treatment of cancer (21). Previously, this goal was approached using a majority-voting scheme to rank genes across various cell types, concentrations, and timepoints. This approach generates a prototype ranked list (PRL) that represents the consistent ranks of genes across several cell lines in response to a specific drug (22).

Once both the disease profile and the disease-specific response to multiple compounds are available, rank-based correlation analysis can be used to quantitatively identify compounds that can revert the transcriptional changes that distinguish diseased samples from healthy ones (5, 9). Nonetheless, monotherapy in cancer is highly susceptible to the development of resistance following an initial response to treatment (23). Combination therapy, or the simultaneous administration of multiple drugs to treat a disease, has evolved into the standard pharmacological regimen for treating complex diseases such as cancer. Combination therapy prevent tumor evolution and help inhibit the development of drug resistance in cancer, thereby improving patient survival (24).

The *in silico* prediction of responses to drug combinations is an active topic of research in computational biology (25–27). Recently, Pickering (2021) found that the transcriptional re-sponse signatures to 856,086 unique two-drug combinations could be predicted based on the 1,309 drug response profiles to compounds tested by the Connectivity Map Build 2 project (parent of the LINCS-L1000 project) (28). After analyzing 148 independent studies involving 257 treatment combinations from the Gene Expression Omnibus (GEO) database, it was found that averaging the expression profiles of individual treatments provides 78.96 percent accuracy in predicting the direction of differential expression for the combined treatment (29). How-ever, different from the Connectivity Map project—where the generation of combinatorial response profiles is possible thanks to the availability of single response profiles for each individual compound (30)—the LINCS-L1000 project does not provide robust single drug response profiles, making the generation of meaningful combinatorial profiles not yet possible.

Here, we present *retriever*, a tool that uses correlation analysis and hierarchical collapsing to extract robust disease-specific transcriptional drug response signature profiles that are consistent across time, concentration, and cell line, from data provided by the LINCS-L1000 project. We integrated these transcriptional drug response profiles with a single-cell RNA-seq signature representing the transcriptional changes observed in triple-negative breast cancer cells compared to healthy breast epithelial cells.

We used these two signatures—the transcriptional changes induced by the disease and the cellular responses to drugs—to prioritize drugs and to predict novel drug combinations suitable for the treatment of triple-negative breast cancer. The transcriptional changes associated with TNBC were computed from a single-cell breast cancer RNA-seq atlas built by us after compiling singlecell RNA-seq data obtained from 36 publicly available healthy breast and breast cancer samples. After systematically testing drug response profiles and combinations at both the population and individual patient levels, we identified QL-XII-47 and GSK-690693 as the most promising combination of kinase inhibitors for treating TNBC. In addition to recommending drug combinations, the profiles returned by *retriever* allow for the characterization of possible mechanisms of action of the identified compound(s) to reverse the disease’s transcriptional profile towards a healthy-like state.

## Methods

### Single-cell RNA-seq datasets

We collected publicly available single-cell RNA-seq count matrices for healthy breast tissues and breast cancer samples from multiple sources (see Data Availability). Each sample’s gene identifiers (IDs) were translated into current gene symbols using the dictionary of gene ID synonyms provided by ENSEMBL (31). Datasets were loaded into R and integrated into a single ‘Seurat’ object (32). Data were then subjected to quality control keeping only cells with a library size of atleast 1,000 counts and within the 95 percent confidence interval of the prediction of the mitochondrial content ratio and detected genes in proportion to the cell’s library size. We also removed all cells that had mitochondrial proportions greater than 10% (33). We then normalized, scaled, and reduced the dimensionality of the data through Principal Component Analysis (PCA) using the default functions and parameters included in Seurat. Data integration was performed using Harmony (34). UMAP projec-tions of the data were extracted using the top 20 dimensions returned by Harmony (35). Cell clustering was performed using the functions included in Seurat for this purpose, with a resolution of 0.01, using UMAP embeddings as the source for the nearest neighbor network construction.

### Cell type assignation

Assignation of cell types was performed based on the expression of the marker genes reported by Wu et al. (36, 37). Epithelial cell subtypes were assigned using the Nebulosa package by querying cells expressing *ESR1, PGR*, and *ERBB2* receptors (38).

### Single-cell RNA-seq differential expression analysis

Transcriptional signatures of triple-negative breast cancer epithelial cells compared to healthy epithelial cells were quantified by differential expression using MAST (39). We compared (Supplementary Fig. S1 and Fig. S2) all epithelial cells from cancer patients (5,730 cells, 31.73%) against those from healthy donors (12,323 cells, 68.26%). In addition, we compared the cluster of patient-derived epithelial cells depleted of hormone and growth factor receptor expression (enrichment for *ESR1*^−^, *PGR*^*−*^, and *ERBB2*^*−*^ cells—2,998 cells, 16.6%) against all healthy epithelial cells (12,323 cells, 68.3%), against the cluster of epithelial cells depleted of hormone and growth factor expression (6,117 cells, 33.9%) derived from healthy individuals, and against the cluster of epithelial cells enriched for triple-positive cells (*ESR1*^+^, *PGR*^+^, and *ERBB2*^+^) from healthy individuals (6,206 cells, 34.4%). We selected the comparison of the cluster enriched with triple-negative cancer cells with the cluster enriched for triple-positive healthy epithelial cells as the disease-specific transcriptional profile, as this profile showed the highest Spearman Correlation Coefficient 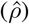 with tissue-level fold-changes computed on TCGA breast cancer data.

### Pseudo-bulk analyses

Pseudo-bulk profiles were computed by summing up all expression values for the same gene across cells of the same individual. We applied this procedure to the triple-negative epithelial cells derived from patients against the triple-positive epithelial cells from healthy individuals. The computed pseudo-bulk profiles from diseased and healthy individuals were then compared using DESeq2 (40).

### TCGA breast cancer dataset analysis

Upper-quantile FPKM-transformed RNA-seq data at the tissue level as well as the associated metadata were downloaded from the TCGA breast cancer (TCGA-BRCA) project using the ‘TCGAbiolinks’ Bioconductor package (41). Samples labeled as ‘healthy’ and ‘basal’ were collected and differential expression between these groups was evaluated using DESeq2 (40).

### Comparisons between data-types

The log_2_ fold-changes of expression between cancer and control samples at the single-cell level, pseudo-bulk level and tissue level were used to validate the computed signatures describing the transcriptional changes in triple-negative breast cancer compared to healthy epithelial cells using the Spearman correlation coefficient (42).

### Application of *retriever* to triple-negative breast cancer

The transcriptional response profiles of triple-negative breast cancer cell lines that were treated with different concentrations and treatment lengths of different compounds available in the LINCS-L1000 project were collected using the ‘ccdata’ Biocon-ductor package, which distributes preprocessed LINCS-L1000 signatures for 1,001 genes (30). The LINCS-L1000 project provides these profiles for three different TNBC cell lines (BT20, HS578T, and MDAMB231). We used these data to compute the disease-specific transcriptional response profiles with *retriever* (default correlation threshold *ρ* = 0.6), grouping profiles by exact matching of their compound, cell line, concentration and time annotations.

### Generation of drug combination transcriptional response profiles

All possible two-drug combinations were generated by averaging the computed disease-specific transcriptional response profiles.

### Drug repurposing analysis

Using the set of overlapping genes between the LINCS-L1000 project and differential expression analysis from single-cell RNA-seq data, we performed Spearman correlation analyses with independent disease-specific drug response profiles and drug combination profiles. We also calculated 95% confidence intervals, p-values, and false-discovery rates (FDR) (43). Compounds were then ranked based on their correlation coefficients, from negative to positive. For individual sample analysis, we compared the cells from each sample to a consistent set of control cells and performed Spearman correlation analyses with both the independent disease-specific drug response profiles and the drug combination profiles.

### Mechanisms of action prediction

To identify possible mechanisms of action for drugs or drug combinations, we took the disease-specific transcriptional response profiles and used these as input for GSEA, with the MSigDB Hallmarks gene set as reference to compute the enrichment of pathways as well as the di-rectionality of the effect (44). Pathways with FDR *<* 0.05 were considered significant. To compare drug-induced and disease-associated changes, GSEA with the Hallmark and GO Biological Process 2023 gene sets (45) was also run on the TNBC signature and on the drug profiles restricted to the 610 genes they share, using fgsea (46).

### Benchmarking and public drug sensitivity data

Using the same LINCS-L1000 TNBC signatures and single-cell TNBC signature, we compared *retriever* with metaLINCS (version 0.9.0) (47), the Prototype Ranked List (PRL), reimplemented following Iorio et al. (22), and naive averaging of all experiments per compound. As ground truth independent of LINCS-L1000, we used the median GDSC1 (release 8.5) area under the dose-response curve (48) across 21 ER-negative/HER2-negative breast cancer cell lines not profiled by LINCS-L1000 (DepMap 22Q2 annotation (49)). Accuracy was the Spearman correlation between predicted reversal scores and sensitivity across the 90 compounds scored by all methods (95% bootstrap confidence intervals), and robustness was the correlation between rankings rebuilt from each pair of TNBC cell lines. GDSC1 sensitivity to QL-XII-47 and GSK-690693 across breast cancer cell lines, and growth-rate metrics (50) for ipatasertib and Torin2 in breast cancer and non-tumorigenic breast cell lines (HMS LINCS dataset 20344), were used to assess the selectivity of the prioritized compounds.

### Cell Lines and Cell Culture

Human breast adenocarcinoma cell lines DU4475 and BT20 were purchased from the American Type Culture Collection (ATCC), while CAL120 was obtained from Leibniz-Institut DSMadeno Z-Deutsche Sammlung von Mikroorganismen und Zellkulturen GmbH. In all cases, the provider supplied an authentication certificate. To ensure the integrity of the cell lines, routine mycoplasma testing was conducted using the Lonza MycoAlertTM kit (LT07-418). None of the cell lines used in this study appeared in the database of commonly misidentified cell lines maintained by the International Cell Line Authentication Committee and NCBI Biosample. The cell lines were not passaged more than 7 times, with the highest passage used being passage 14. DU4475 cell line was cultured in RPMI-1640 media (ThermoFisher Cat:A1049101) supplemented with 10% Fetal Bovine Serum (FBS; ThermoFisher Cat:10500064) and 1% penicillin–streptomycin (ThermoFisher; 10,000 U/mL). CAL120 cell line was maintained in DMEM-High glucose media (ThermoFisher Cat:11965084) with the same supplements, while BT20 cell line was cultured in EMEM (ThermoFisher Cat:31095029) media with identical supplementation.

### Cell Viability Assays

BT20 and CAL120 cells were initially seeded at a density of 4000 and 2500 cells/well, respectively, in a 96-well plate and allowed to incubate overnight. After 24 hours, the culture media was replaced with media containing predetermined concentrations of selected drugs, namely BMX/BTK Inhibitor II, QL-XII-47 (CAS 1469988-75-7, Sigma Cat:5307580001), and GSK-690693 (Sigma Cat:SML0428-5MG), and incubated for an additional 72 hours. For the suspended cell line DU4475, cells were not counted before seeding, and drugs were added from the initiation of culturing. Cell viability for the drug assay was evaluated using the ATP-based CellTiter-Glo® 3D Cell Viability Assay kit (Promega Cat:7570). The luminescence signal of each well was measured using a bottom-reading luminescent plate reader (SpectraMax M5 Multimode microplate reader). In each independent experiment, a minimum of three technical replicates were employed, and at least three independent experiments were conducted. The viability expected under Bliss independence was computed as the product of the mean fractional viabilities of the single agents (51), with bootstrap confidence intervals for the difference between observed and expected viability. The combination was compared with each single agent using two-sided Welch *t*-tests, with P values adjusted across cell lines using Holm’s method (52).

## Results

### The *retriever* algorithm

The *retriever* algorithm extracts disease-specific drug response profiles using three steps (Figure 1). The first step summarizes the cellular responses at different time points after the application of the drug, the second step summarizes the responses at different concentrations, and the third step summarizes the responses across different cell lines. This provides robust, disease-specific transcriptional response profiles based on the responses observed in all cell lines used as surrogates of a specific disease.

**Fig. 1.**
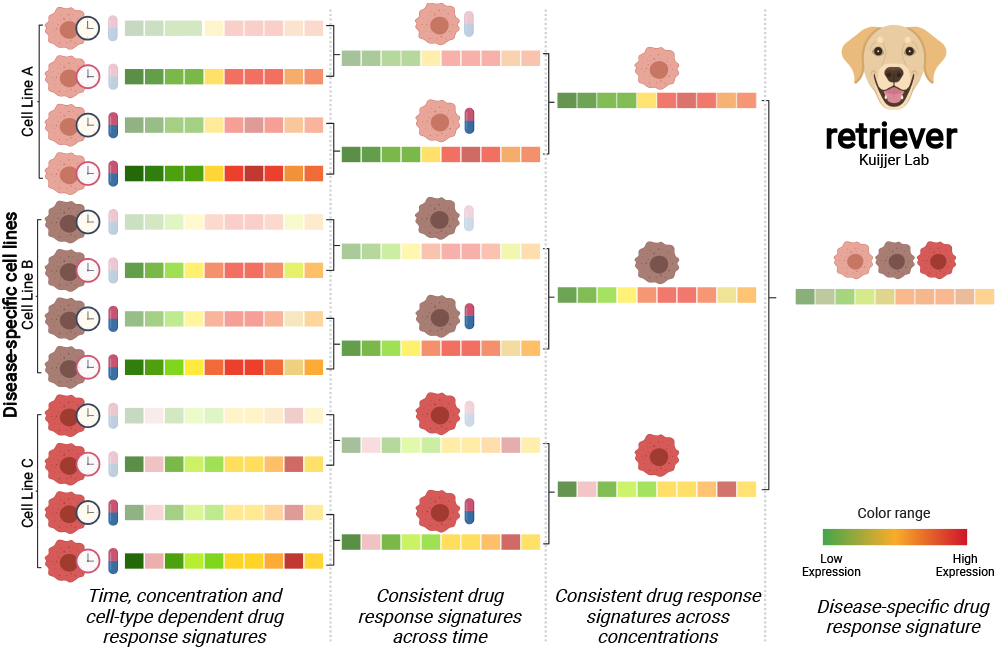
Overview of the *retriever* algorithm. *retriever* generates disease-specific transcriptional drug response signatures by merging transcriptional signatures over time, concentration, and cell-type. These signatures can then be matched to single-cell or bulk expression profiles to predict drugs and drug combinations most likely to be effective in treating a disease.

In the first step, *retriever* takes the response profiles of a given cell line to the same compound under the same concentration, but at different time points, and averages these. Then, the descriptive power of the extracted profile to represent the transcriptional response to a drug at a given concentration in the cell line, consistent across time, is evaluated using Spearman’s correlation coefficient.

The averaged profile is returned if the correlation with the computed profile is larger than a user-defined threshold (default *ρ* = 0.6). This threshold can take values between 0 and 1, representing the percentage of agreement between the cellular responses to the same compound. If the threshold is too close to one, very few averaged profiles will be returned due to strong filtering, and if it is defined to close to zero, very noisy profiles (similar to simply averaging all the available profiles) will be returned. The original drug response profiles that do not reach this threshold are removed. The averaged profile is then recomputed using all response profiles that reached the threshold. This procedure ensures the removal of aberrant or insufficient cellular responses. Only averaged signatures of at least two profiles are used in the second step.

In the second step, *retriever* takes the stable time-consistent signature profiles of a compound at a particular concentration in the same cell line. To summarize the response at different concentrations, it applies the same procedure described in the first step over the averaged profiles. The profiles returned by the second step are stable transcriptional response profiles that are consistent across time and drug concentration in a specific cell line.

The last step extracts disease-specific drug response profiles by again applying the procedure described in step 1 to the stable response profiles to the compound in all disease-specific cell lines available in the LINCS-L1000 project. The profiles returned by the third step are robust disease-specific response signatures representing transcriptional changes to a specific compound.

### Single-cell RNA-seq atlas of breast samples

To showcase how *retriever* can be used to prioritize drugs, we applied it to single-cell data from breast cancer and healthy breast tissues. For this, we compiled 36 publicly available single-cell RNA-seq count matrices from breast samples (26 diseased and 10 healthy). In total, we combined 109,097 cells into a single Seurat object, maintaining the sample of origin metadata. Following quality control (see Methods section), 77,384 cells were retained for further analysis (Figure 2A), of which 30,790 were derived from healthy (left panel in Figure 2B) and 46,594 from cancer samples (middle panel in Figure 2B). Cell types were assigned to the nine identified clusters in the low dimensional representation (Figures 2C, S3) using markers reported by Wu et al. (36, 37).

**Fig. 2.**
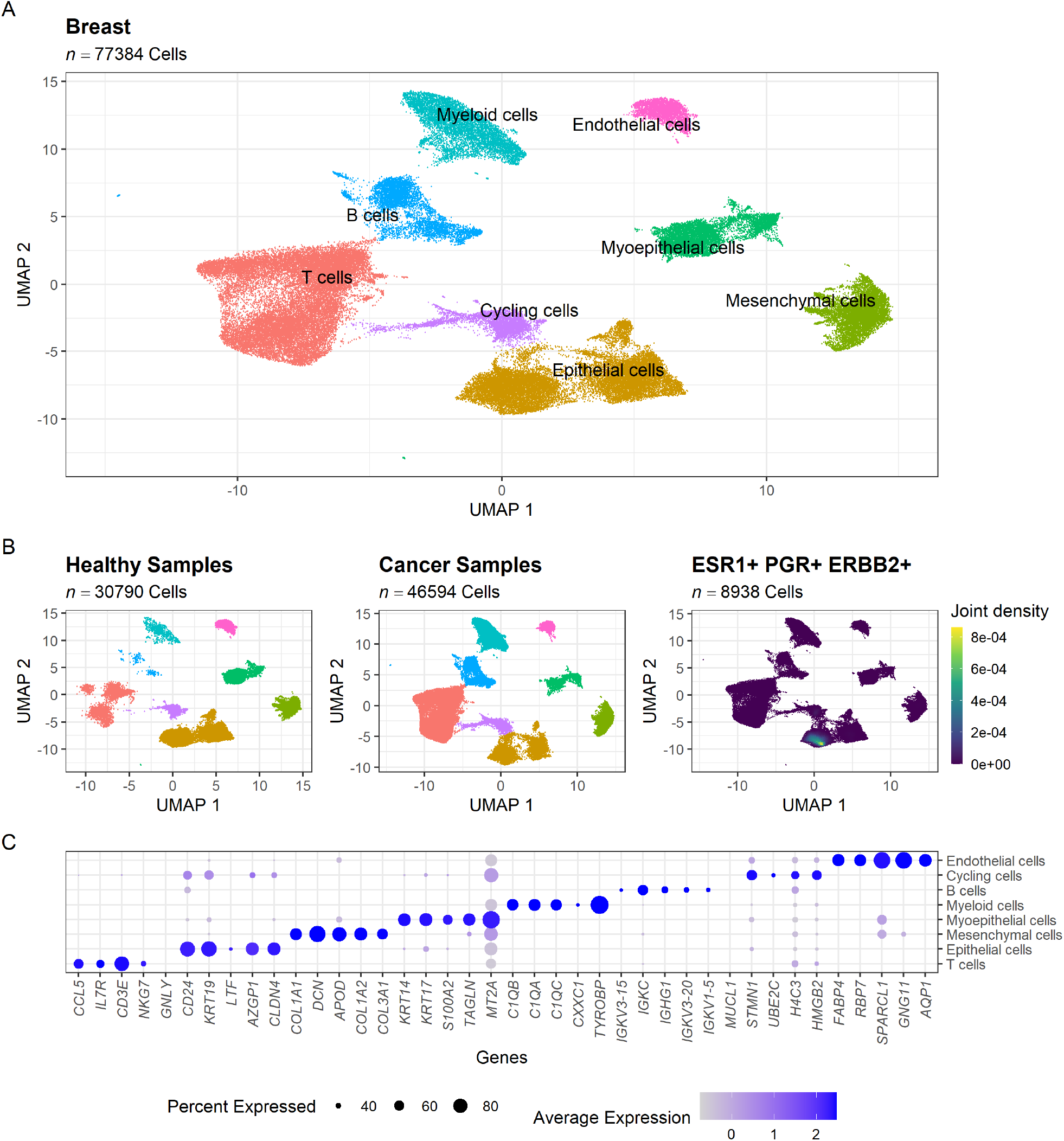
Single-cell atlas of breast samples. **(A)** UMAP projection of the integrated 77,384 cells from 36 breast samples. **(B)** UMAP projection of the healthy (left) and cancer (center) cells, and visualization of triple-positive epithelial cells in the cancer samples. **(C)** Dotplot representing the normalized expression level and percentage of cells expressing the top five differentially expressed genes for each cell type.

### Transcriptional signatures associated with triple-negative breast cancer at the single-cell level

To identify the transcriptional signature that best represents the changes associated with triple-negative breast cancer, we first identified the single cells with a triple-negative phenotype within the epithelial cluster of cells. To do so, we queried cells expressing ESR1, PGR, and ERBB2 receptors, which allowed us to assign a cluster of cells enriched for the expression of the three receptors (Right panel in Figure 2B); cells belonging to the cluster enriched for triplepositive cells will be further referred as “triple-positive-like” cells in the remainder of this work. While cells belonging to the other cluster will be referred to as “triple-negative-like” cells.

We compared the triple-negative-like breast cancer cells with different healthy epithelial cell subpopulations, including all epithelial cells, cells belonging to the triple-negative cluster, and cells belonging to the triple-positive cluster of epithelial cells. We then compared fold changes to those derived from the population level-data from TCGA (see Methods section). The most representative profile was the one comparing triple-negative-like epithelial cancer cells with healthy triple-positive-like epithelial cells. Note that, while approximately half of all healthy epithelial cells are assigned to the triple-positive cluster, these receptors are expressed in only 20% of these cells, and at lower levels compared to breast cancer cells (Supplementary Fig. S1).

As a result, we used MAST to perform differential expression analysis at the single cell level between triple-negative-like epithelial cancer cells (*n* = 2,998) and healthy triple-positive-like epithelial cells (*n* = 6,206), identifying 205 differentially expressed genes (106 upregulated and 99 downregulated) with absolute log_2_ fold-changes larger than 1 and false discovery rate lower than 0.05 (Figure 3A and Supplementary Table S9). Downregulated genes included estrogen-responsive and luminal genes (*TFF1, AGR2, PIP, STC2, AREG*) and NF-*κ*B targets (*CXCL8, CXCL1, IER3*), consistent with the loss of hormone receptor signaling that defines TNBC, whereas upregulated genes included basal keratins (*KRT14, KRT15*), *VIM, S100A8*, and *S100A9*, reflecting the basal-like phenotype of TNBC.

**Fig. 3.**
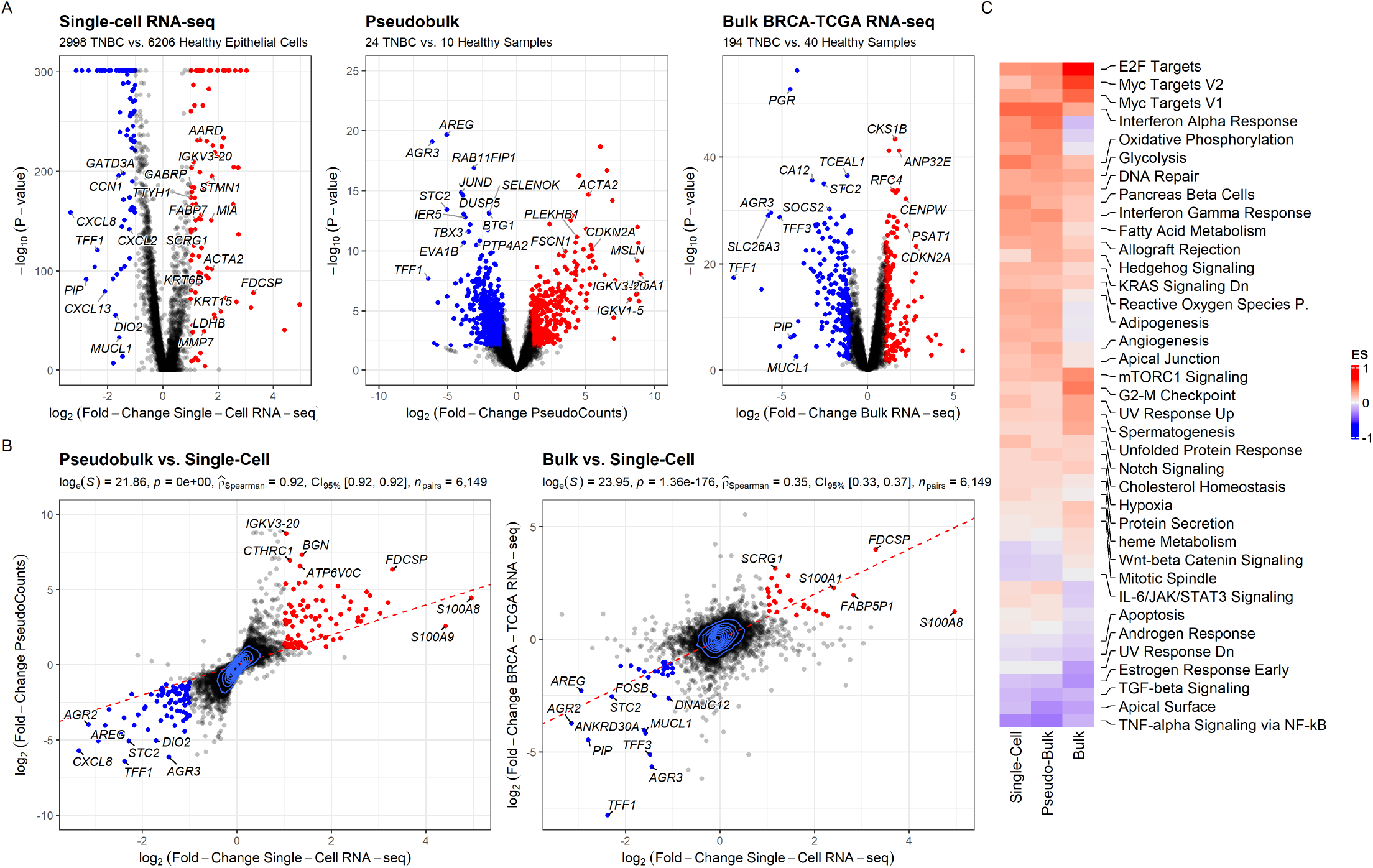
Transcriptional changes identified in TNBC cells. **(A)** Volcano plots report the difference between TNBC and healthy samples, on the left based on single-cell RNA-seq count data computed using MAST, in the middle using the pseudo bulk measures in each single-cell RNA-seq sample, and on the right using the bulk RNA-seq data from the BRCA-TCGA project. Each dot represents a gene. Dots are color coded, in red if the log_2_ fold-change is larger than 1 (upregulated in TNBC; 106 genes in the single-cell comparison) and in blue if the log_2_ fold-change is smaller than −1 (downregulated in TNBC; 99 genes). **(B)** Comparisons of the transcriptional changes associated with TNBC at the single-cell, pseudo bulk and tissue level. Each dot represents a gene. Dots are color coded as in Figure 2A. **(C)** Single-sample GSEA Enrichment Score (ES) for the transcriptional changes between TNBC and healthy cells at different levels of resolution. Labeled pathways are those that show same trend (positive or negative ES) in the three data types.

After testing using both the hypergeometric test through Enrichr (45) and gene set enrichment analysis (GSEA) using the Hall-mark MSigDB signatures as reference gene sets (44), we found that the differentially expressed genes were associated with the activation of *Oxidative Phosphorylation Pathway, Interferon Alpha Response*, as well as *Myc Targets*, and the downregu-lation of *TNFα Signaling via NF-κB, Estrogen Response, UV Response Up, Apoptosis, Hypoxia, Unfolded Protein Response, IL-6/JAK/STAT3 Signaling, Inflammatory Response*, and the *Androgen Response* pathway (Figure 3C, Supplementary Table S1). Many of these pathways are known to be highly associated with TNBC molecular phenotype. For example, activation of both the oxidative phosphorylation pathway and of Myc targets is associated with a worse outcome of the disease (53, 54), and the crosstalk between the interferon alpha response pathway and NF-*κ*B is associated with drug resistance and tumor progression in this malignancy (55). The downregulation of the estrogen response pathway as well as apoptosis and hypoxia are markers of TNBC (56). In addition, downregulation of androgen and inflammatory responses is associated with worse prognosis and higher chemotherapy responsiveness respectively (57, 58).

We found 6,149 genes expressed in both the single-cell RNA-seq datasets and the bulk RNA-seq from the TCGA-BRCA project. Spearman correlation coefficients between the expression changes computed at the single-cell level (Left panel in Figure 3A), pseudo-bulk level (Middle panel in Figure 3A) and tissue level (Right panel in Figure 3A), revealed a monotonic positive association between these different levels (Figure 3B), supporting the descriptive power of the computed differential single-cell expression signature to describe transcriptional changes associated with TNBC, and ensuring that the selected cell type as well as the pseudo-bulk samples are high-resolution descriptors of the diseased tissues.

### Applying *retriever* to extract TNBC-specific transcriptional drug-response signatures

We collected 4,899 response profiles measured in TNBC cell lines from the LINCS-L1000 project using the ‘ccdata’ package available in Bioconductor (30). These profiles correspond to the expression changes of 1,001 genes in response to 205 perturbagens (176 small molecules, most of them kinase inhibitors, and 29 ligands) in three TNBC cell lines (BT20, HS578T, and MDAMB231). Small molecules were profiled at four concentrations (0.08, 0.4, 2, and 10 *µ*M) and two time points (6 and 24 hours).

An illustration of the step-by-step process of constructing a gen-eralized response profile to a compound across three TNBC cell lines is presented in Figure 4 using the QL-XII-47 compound as a case example.

**Fig. 4.**
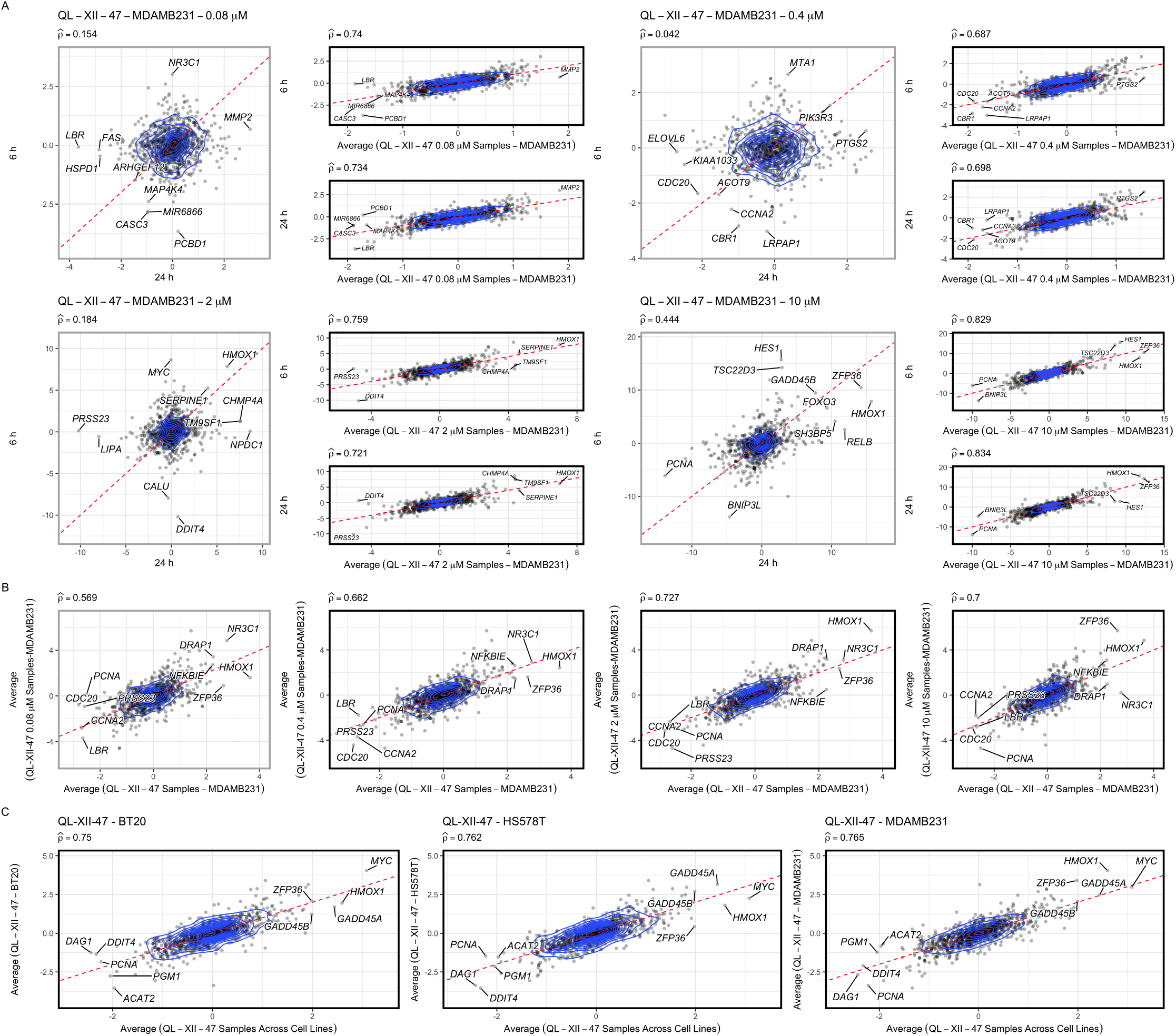
Case example showing the construction of a single transcriptional response profile to a compound across three TNBC cell lines. Frames of the scatterplots displaying the relationship between the profiles and the averaged profiles are color coded, in black if the computed Spearman correlation coefficient 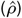 is larger than 0.6 or in gray otherwise. **(A)** Generation of a time-consistent response profile. **(B)** Generation of a time and concentration-consistent response profile. **(C)** Generation of a time, concentration, and cell-line-consistent response profile.

#### 1. Computing time-consistent generalized response profiles

In the first step (Figure 4A), we computed time-consistent generalized response profiles. To achieve this, we took the response profile of the MDAMB231 cell line to QL-XII-47 under the same concentration at two different time points (6 and 24 hours) and averaged them. When we compared the transcriptomes of the cell line exposed at two different time points, we found little or no correlation (large boxes labeled in gray) for each concentration (0.08*µ* M, 0.4*µ* M, 2*µ* M, and 10*µ* M). However, we found that their averaged profile displays high predictive power (Spearman Correlation Coefficient 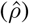 > 0.65 in all cases) of the cellular response to the compound, independent of the time of treat-ment. When present, we removed the profiles that did not correlate with the averaged profile above 0.6 (A total of 11.33% of the profiles). This procedure allows to identify concentrations in which the compounds induces a differ-ent, aberrant or insufficient cellular responses to the drug and also allows us to compute a single robust response profile to the compound.

#### 2. Computing time- and concentration-consistent response profiles

In the second step (Figure 4B), we extract time and concentration-consistent responses. For this purpose, we averaged the time-consistent profiles computed in the first step. As before, we removed the profiles that did not correlate with the averaged profile above 0.6, such as the profile present in the first panel of Figure 4B, which showed a different cellular response at the concentration (A total of 7.62% of the profiles computed in Step 1). We then recomputed the profile for the compound in the MDAMB231 cell line with all profiles that had correlation coefficients above the threshold.

#### 3. Generating disease-specific drug response profiles

In the last step (Figure 4C), we processed the outputs of the second step to extract a generalized response profile to a compound across the three TNBC cell lines. To do this, we performed independently steps 1 and 2 for QL-XII-47 in the other two TNBC cell lines available in the LINCS-L1000 project (BT20, HS578T) and averaged these profiles.

After summarizing the time points, concentrations, and cell lines for all the 205 compounds tested in TNBC cell lines, we ended with robust transcriptional response profiles of 151 (73.66%) compounds (Supplementary Table S2) that are specific for TNBC. The remaining 54 compounds were removed, as they did not show a stable response across time points, concentrations, and cell lines.

Having a single generalized disease-specific response profile for each compound allowed us to (1) generate compound combinations *in-silico* as candidates for the treatment of TNBC (Supplementary Table S3), (2) to rank compounds and compound combinations based on their predicted potential to reverse the disease state towards a healthy-like state (Supplementary Tables S4 and S5), and to (3) interrogate the computed profiles using rank-based gene set enrichment analysis (GSEA) to predict possible mechanisms of action of the compounds when used to treat TNBC.

Our method to extract disease-specific drug response profiles is implemented in the ‘*retriever*’ R package (see Data Availability section) and allows the computation of disease-specific transcriptional drug response signatures for different cancer types available in the LINCS-L1000 project.

### Ranking compound candidates for the treatment of TNBC

We used the 151 TNBC-specific transcriptional response profiles extracted with *retriever* to compute 11,325 response profiles to nonredundant drug combinations. To do so, we calculated the averaged effect sizes of Spearman correlation between the response profiles and the differential single-cell transcriptional signatures and then ranked the compounds and compound combinations likely to specifically antagonize the TNBC-specific transcriptional signatures. Since Spearman correlation is a rank-based approach, this method can quantitatively identify inverse effects induced by drugs using the response profiles and disease-associated transcriptional signatures. Thus, we computed the Spearman coefficients, 95% confidence intervals, and p-values corrected for multiple testing (FDR) for all 11,476 profiles (Supplementary Tables S4 and S5). We then ranked the drug combinations by their correlation coefficients, from negative to positive. A negative value represents the expected inverse effect of the drug against the expression changes observed in the disease towards a healthy-like state.

We found that QL-XII-47 is the most promising compound for reversing the transcriptional profile of TNBC back to a healthy-like state. Followed by Torin-2, Torin-1, QL-X-138, and WYE-125132. This top candidate remains consistent even when evaluating each disease signature for individual diseased samples included in the generated atlas. Specifically, QL-XII-47 emerged as the top candidate in 12 of the 25 (*P* = 7.12 *×* 10^−27^; Top panel in Figure 6) analyzed samples (one sample was excluded due to insufficient cells for differential expression analysis after quality control).

QL-XII-47 is a highly effective and selective Bruton tyrosine kinase (BTK) inhibitor that covalently modifies Cys481 of the protein. QL-XII-47 has an IC_50_ of 7 nM for inhibiting BTK kinase activity and induces a G1 cell cycle arrest in Ramos cells (B lymphocytes from Burkitt lymphoma), which is associated with significant degradation of the BTK protein. It was also shown that, at sub micromolar concentrations, QL-XII-47 inhibits the proliferation of B-cell lymphoma cell lines (59). Furthermore, other BTK inhibitors have shown to reduce TNBC cell viability (60). In addition, independent validation of sensitivity to QL-XII-47 has been tested before in two other TNBC cell lines (HCC70 and HCC1806) that were not included in the LINCS-L1000 project. Low toxicity (lower than Torin-2) and more than 50% reduction of the growth rate (GR_max_) was found for both cell lines (61).

Through enrichment analysis, we found that QL-XII-47 may act in TNBC through activation of the *TNF-alpha Signaling via NF-κB, Hypoxia-associated genes*, the *Inflammatory Response*, the *IL-2/STAT5 Signaling pathway*, and by deactivating genes involved in *Fatty Acid Metabolism* (FDR *<* 0.05 in all cases, Fig 5B).

**Fig. 5.**
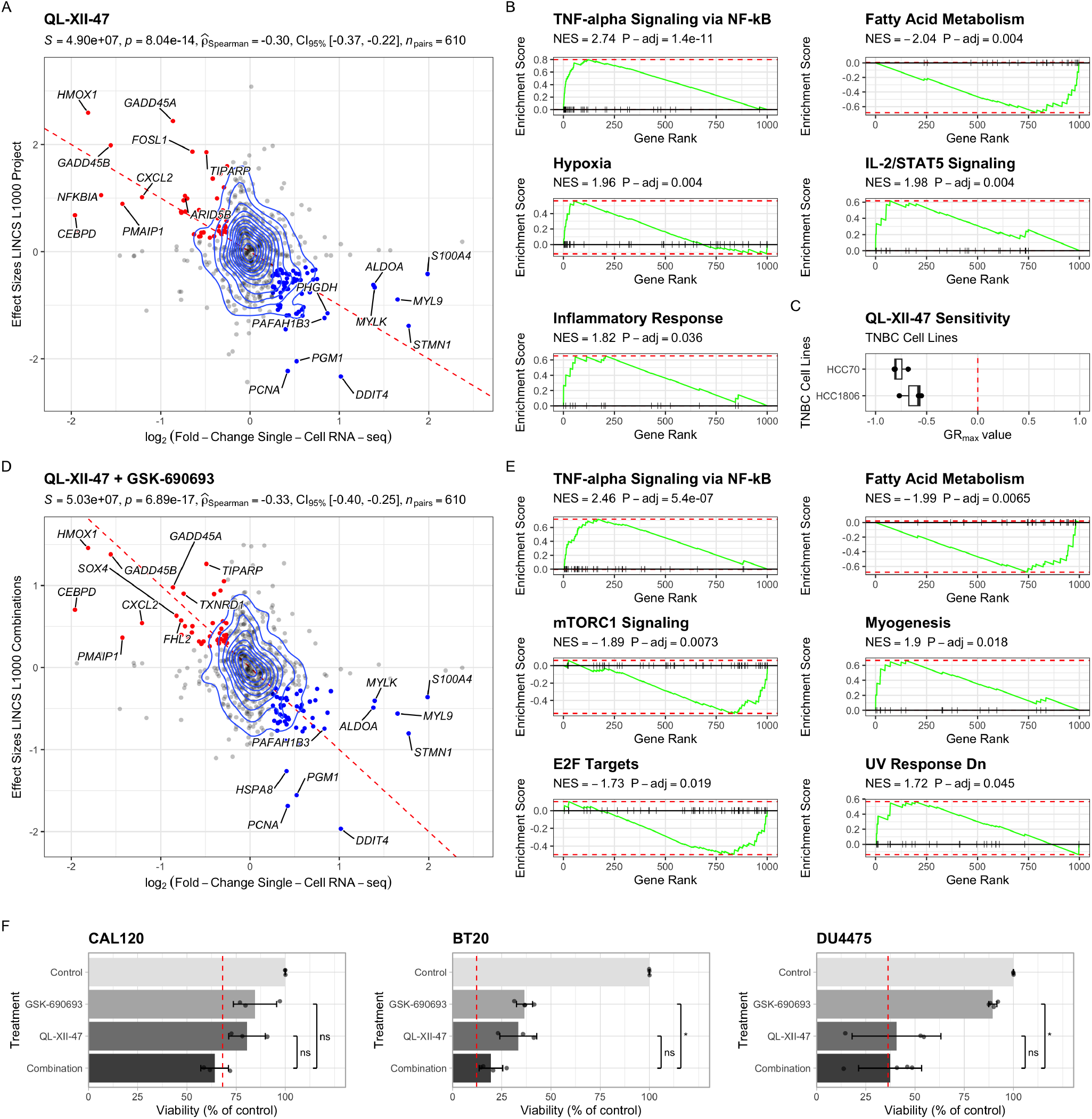
Correlation analysis and mechanism of action prediction for the disease-specific drug response profiles in TNBC. **(A)** Spearman correlation analysis between the expression changes associated with TNBC and the disease-specific drug response signature of QL-XII-47. Each dot represents a gene. Dots are color coded, in red if they are expected to be upregulated by the drug, in blue if they are expected to be downregulated by the drug, and in gray if no significant change is expected. The red line represents the perfect match between both profiles. Density lines reflect the number of dots in each section of the plot. **(B)** Mechanisms of action predicted for QL-XII-47 in TNBC based on the enrichment of biological pathways in the disease-specific transcriptional drug response signature. **(C)** Independent sensitivity evaluation of the effect of QL-XII-47 in two other TNBC cell lines. **(D)** Spearman correlation analysis between the expression changes associated with TNBC and the combination signature of QL-XII-47 and GSK-690693. Each dot represents a gene. Dots are color coded as in (A). The red line represents the perfect match between both profiles. Density lines reflect the number of dots in each section of the plot. **(E)** Mechanisms of action predicted for the mixture of QL-XII-47 and GSK-690693 based on the enrichment of their combination drug response signature. **(F)** Comparisons of cellular viability among three distinct breast cancer cell lines following treatment with 0.6 *µ*M QL-XII-47, 0.8 *µ*M GSK-690693, and the combination of both compounds under same concentration. Bars show mean *±* SD and dots show individual replicates. The red dashed line shows the viability expected under Bliss independence. The combination was compared with each single agent using two-sided Welch *t*-tests with Holm adjustment across cell lines: ns, P *>* 0.05; *P ≤ 0.05.

### Ranking compound combination candidates for the treatment of TNBC

After testing combinations of disease-specific drug response profiles, we found that the combination of QL-XII-47 with GSK-690693 – a pan-AKT kinase inhibitor that reduces tumor cell proliferation and induces tumor cell apoptosis (62, 63)– was the best-performing drug combination to revert TNBC signatures. This combination consistently ranked as the best across individual diseased samples in our atlas. Specifically, QL-XII-47 and GSK-690693 emerged as the top combination in 5 of the 25 analyzed samples (*P* = 5.37 *×*10−21; see bottom panel in Figure 6).

**Fig. 6.**
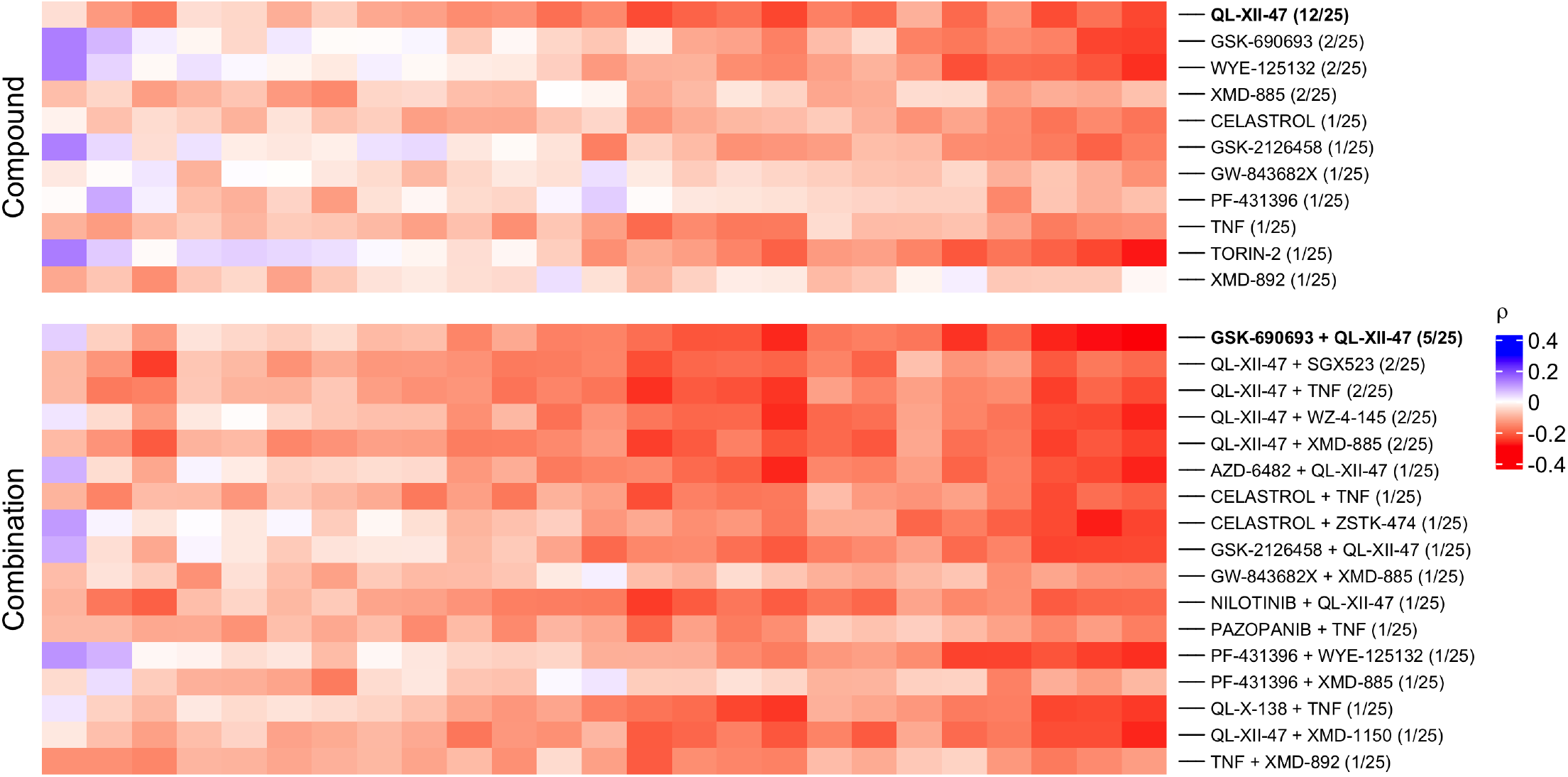
Heatmap displaying the Spearman correlation coefficients for the top-ranking compounds and compound combinations in each sample within the generated atlas. The compounds or combinations that ranked 1^st^ in at least one sample are included. The number of times each compound or combination ranked 1^st^ across all samples is indicated in parentheses.

Gene set enrichment analysis predicted that this combination may act through the activation of *TNF-alpha signaling via NF-κB, Myogenesis, Apoptosis*, and *UV Response Dn*, and the deactivation of *Fatty Acid Metabolism, mTORC1 Signaling*, and *E2F targets* (FDR *<* 0.05 in all cases, Fig 5E). These pathways have previously been associated with the inhibition of triple-negative breast cancer growth and metastasis (64–66), and thus highlight the potential of our computational approach to prioritize drugs by integrating cell-type specific profiles obtained from single-cell RNA-seq data sets with disease-specific transcriptional drug response profiles. On the genes shared with the TNBC signature, the combination reversed the two Hallmark pathways most strongly altered in TNBC cells, *TNF-alpha Signaling via NF-κB* (NES −2.4 in TNBC and +2.3 for the combination) and *Apoptosis* (−1.9 and +1.9; FDR *<* 0.05 in all cases), and 15 of the 16 GO Biological Process terms enriched for the combination had the opposite sign in the TNBC signature, although none of these was individually significant in the disease (Supplementary Fig. S6).

Moreover, we performed experimental validation of the identified top combination in three different breast cancer cell lines (CAL120, BT20, and DU4475). We compared the cellular viability (Supplementary Table S6) of the cells under baseline conditions and after treatment with 0.6 *µ*M QL-XII-47, 0.8 *µ*M GSK-690693, and the combination of both compounds under the same concentration. QL-XII-47, the top-ranked compound, reduced viability in all three cell lines, to 33.4% in BT20, 40.6% in DU4475, and 80.5% in CAL120 (Figure 5F). The combination produced effects consistent with Bliss additivity in CAL120 and DU4475 and below additivity in BT20, and did not differ significantly from QL-XII-47 alone after correction for multiple testing. This is consistent with how *retriever* builds combination profiles, as the mean of two single-drug signatures, which assumes independent drug action and therefore does not predict synergy. DU4475 replicates were highly variable (coefficient of variation 55.6% for QL-XII-47), probably because this suspension line was not counted before seeding.

Public data support these observations (Supplementary Fig. S5). In GDSC1, BT20 (IC_50_ 0.80 *µ*M) and DU4475 (1.11 *µ*M) were sensitive to QL-XII-47 and CAL120 was comparatively resistant (7.36 *µ*M), mirroring our measurements, and the concentrations used here are below the GDSC1 IC_50_ of QL-XII-47 in all 23 ER-negative/HER2-negative breast cancer cell lines screened. Neither QL-XII-47 nor GSK-690693 was more potent in TNBC than in non-TNBC breast cancer cell lines, whereas the pan-AKT inhibitor ipatasertib and Torin2 were less active in the non-tumorigenic MCF10A and hTERT-HME1 cells than in TNBC cell lines. Thus, *retriever* prioritizes compounds that reverse the TNBC transcriptional state, which does not by itself imply selective toxicity to TNBC cells.

### Benchmarking against existing methods

Compared with metaLINCS (47), the PRL (22), and naive averaging, *retriever* performed similarly (Supplementary Fig. S4). The rankings of all methods correlated with GDSC1 drug sensitivity in 21 TNBC cell lines not profiled by LINCS-L1000 (*retriever ρ* = 0.31, 95% CI 0.08 to 0.51; metaLINCS 0.29, 0.05 to 0.49; PRL 0.34, 0.13 to 0.52; naive averaging 0.28, 0.04 to 0.48), and were similarly ro-bust when one cell line was removed (mean correlation between rankings 0.83 to 0.87). All methods ranked QL-XII-47 among the top six compounds. The added value of *retriever* is there-fore not higher ranking accuracy but the consensus signature it returns for each compound, which enables combination profiles and pathway-level analyses.

## Discussion

Single-cell RNA-seq provides an unprecedented resolution to characterize cellular heterogeneity in cancer. Compared with cell lines, the cells from tumors characterized by single-cell RNA-seq do not exhibit metabolic adaptation due to culturing. In advantage to bulk RNA-seq, single-cell RNA-seq can provide a more accurate signature of the changes observed in specific cells relevant to the disease under study(14, 67). Thus, leveraging the information provided by this new technology is important to accelerate the identification of new personalized treatments that target specific subpopulations of cells in a tissue (68).

However, just as important as characterizing the signatures of the disease in affected cells at the highest possible resolution is to characterize the response profiles of drugs that may help to reverse disease-associated transcriptional changes towards a healthy state. Large consortia including the LINCS-L1000 project provide transcriptional phenotypes at different time points after applying hundreds of compounds to multiple types of cell lines at different concentrations. Even though this information is valuable, its direct application to pathophysiological scenarios is limited due to the difficulty of extracting consistent response profiles in these large data sets (69).

Here, we introduced *retriever*, a tool to extract robust disease-specific drug response profiles from the LINCS-L1000 project. *retriever* mines the information associated with each cell line in the project to select subsets of cell lines that are surrogates of a defined disease, after which it extracts disease-specific response profiles. By doing this, *retriever* maximizes the robustness of the transcriptional signatures that are used to rank candidate compounds for the identification of new treatment strategies. With *retriever*, we hypothesize that if a signature is robust across multiple cell lines representing the same disease, it will likely also be more robust across individuals, as cell lines exhibit genetic and metabolic differences.

We showcased *retriever*’s strength by retrieving time-, dose-, and cell-type consistent transcriptional drug response signatures of TNBC cells from the LINCS-L1000 project and combining it with single-cell RNA transcriptome profiles obtained from a large single-cell breast cancer atlas, which we used to predict novel drug combinations against TNBC. This identified a combi-nation of two kinase inhibitors predicted to be effective together acting on important biological pathways in breast cancer, and thereby antagonize the TNBC-specific signature.

The prediction of drug combinations with *retriever* that we present here, although it is relatively simple and relies on the hypothesis of independent mechanisms of action for each compound, has been established as accurate (predicting the right directionality of the change) in analyses performed on profiles from the Connectivity Map project (30). Nevertheless, our computational approach only allows us to rank compounds and compounds combinations that are suitable for the development of treatments based on the negative correlation of the drug response profiles and the disease signatures. Other promising compounds and drug combinations still need to be validated experimentally to define the right concentration and evaluate if they indeed exhibit a synergistic mechanism of action. Their toxicity across cell lines and, after preclinical tests, potential adverse events in *in-vivo* models and cancer patients also need to be evaluated. Our experimental support is limited to single-dose *in vitro* assays in three cell lines, complemented by public drug sensitivity data, and QL-XII-47 has not been tested clinically. Beyond general transcriptional regulation (11 of the 16 terms), the GO Biological Process terms enriched for the combination point to three processes relevant to TNBC treatment (Supplementary Fig. S6): negative regulation of proliferation, through *CDKN1A* and *CDKN1B*, the expected consequence of AKT inhibition; regulation of apoptosis, through *GADD45A* and *GADD45B*, which are downregulated in TNBC, a disease treated mainly with cytotoxic chemotherapy; and positive regulation of kinase activity, through the receptor tyrosine kinases *ERBB3* and *IGF1R*. Induction of these receptors is consistent with their known feedback activation after AKT inhibition (70), which may limit AKT-based therapies: AKT inhibitors combined with paclitaxel improved progression-free survival in phase II trials in metastatic TNBC (71, 72), but this was not confirmed in phase III (73, 74), suggesting that blocking this feedback may be needed to improve the efficacy of AKT-based combinations.

Thanks to the multiple cell lines available in the LINCS-L1000 project, our approach can be replicated in at least 13 other cancer types, including: *adult acute monocytic leukemia, adult acute myeloid leukemia, cecum adenocarcinoma, colon (adeno)carcinoma, endometrial adenocarcinoma, lung adenocarcinoma, large cell lung carcinoma, small cell lung carcinoma, melanoma, ovarian mucinous adenocarcinoma, prostate carcinoma*, and *triple-negative breast cancer* among other variants (such as local or metastatic tumors for some cancer types), as soon as single-cell RNA-seq data for those cancer types and healthy tissues become available. Considering the increase in single-cell RNA-seq studies being published since the development of the technique, we expect this to be possible for most cancer types represented in the LINCS-L1000 project in the near future (75).

We want to highlight the important ethical considerations involved in using patient-derived data for drug development and repurposing, particularly around data privacy, informed consent, and the reliability of predictive models. To protect patient privacy, it is crucial to adhere to data protection laws, such as HIPAA and GDPR, and to rigorously de-identify data to minimize the risk of re-identification (76). Additionally, datasets must be diverse and representative to prevent bias, ensuring that predictive models are applicable to a broad population. Computational models, including *retriever*, should undergo extensive validation before being used in clinical settings to ensure their accuracy and transparency. Ethical protocols for data sharing must also be established to respect patient autonomy and control over their data. Furthermore, continuous monitoring and validation of drug predictions are necessary to ensure treatment safety, effectiveness, and fairness.

Although *retriever* represents a significant advancement in extracting disease-specific drug response profiles from the LINCS-L1000 dataset, several limitations must be considered when interpreting its results. One key limitation is the restricted scope of gene expression data in the LINCS-L1000 project, which includes expression profiles for only 1,000 genes. While this gene set provides valuable insights into broad transcriptional changes, it may not fully capture the complexity of cellular responses to drug treatments. A possible solution to this limitation relies on imputation techniques to address the missing quantification in the gene expression matrix. The accuracy of imputed values is dependent on the quality of the imputation model and the completeness of the available data. Consequently, there is an inherent risk that the imputed values may not accurately represent the true and complete underlying biological response.

*retriever* also has limitations of its own. Its consistency filter depends on a fixed correlation threshold: 205, 200, 151, 81, and 6 TNBC compounds pass at thresholds of 0.4, 0.5, 0.6, 0.7, and 0.8, and GSK-690693 is removed at 0.7, so the threshold should be treated as a tunable parameter. The filter favors compounds with broad, consistent responses and discards compounds whose effect is specific to one cell line or dose. Consensus profiles are unweighted averages that do not model dose-response relationships, and combination profiles assume independent drug action, so *retriever* cannot predict synergy or recommend concentrations. A negative correlation with the disease signature indicates transcriptional reversal towards the chosen healthy reference, not cytotoxicity or tumor selectivity. Finally, the profiles depend on the cell lines available as disease surrogates; for TNBC, two of the three (HS578T and MDAMB231) are mesenchymal-like, and the compounds profiled in them are mainly kinase inhibitors.

Finally, we have shown that the approach implemented in *retriever* method can predict effective drug combinations for patients with triple-negative breast cancer (TNBC), but its potential goes beyond that. It can also be applied to single-cell RNA sequencing data from individual tumors and other diseases for which a the single-cell transcriptomic profile of a normal control population is available, opening up the road for applications in precision medicine. In line with this, the LINCS project has released datasets for iPSC-derived cardiomyocytes and motor neurons, opening up new possibilities for precision medicine not only in cancer but also in a variety of other diseases (77, 78). By predicting the most effective drug and combination treatments for each patient, clinical trials can be designed to target the right populations with the responsive transcriptional phenotype, leading to more successful outcomes.

## Data Availability

All the data and code required to replicate the analysis as well as the figures and tables are available at https://github.com/dosorio/L1000-TNBC, including the code to download the public data (GDSC1, DepMap, HMS LINCS) used in the benchmark. The *retriever* package is available from the Kuijjer Lab repository https://github.com/kuijjerlab/retriever or from the CRAN repositories https://cran.r-project.org/package=retriever, and it is implemented as an R multiplatform package that can run on standard laptops or desktops with around 16 GB of RAM, making it accessible for most users. It is designed to work on Windows, macOS, and Linux. While the package can function with modest hardware, performance may vary based on dataset size and complexity. For larger datasets, systems with more RAM or cloud-based resources may improve efficiency.

The following datasets were used to construct the single-cell RNA-seq breast atlas used in this study:

1. Tabula sapiens wild-type mammary gland. Data from (79) Quake, Stephen R., and Tabula Sapiens Consortium. *“The Tabula Sapiens: a single cell transcriptomic atlas of multiple organs from individual human donors*.*”* bioRxiv (2021). Accessed through: https://tabula-sapiens-portal.ds.czbiohub.org/
2. Wild-type data from seven individuals reported by (80) Nguyen, Quy H., *et al. “Profiling human breast epithelial cells using single cell RNA sequencing identifies cell diversity*.*”* Nature communications 9.1 (2018): 1-12. GEO accession code: GSE113197.
3. Wild-type data from five individuals reported by (81) Bhat-Nakshatri, Poornima, *et al. “A single-cell atlas of the healthy breast tissues reveals clinically relevant clusters of breast epithelial cells*.*”* Cell Reports Medicine 2.3 (2021): 10021955. GEO accession code: GSE164898.
4. Triple-negative breast cancer data from 9 individuals from (36) Wu SZ, Al-Eryani G, Roden DL, Junankar S *et al. A single-cell and spatially resolved atlas of human breast cancers*. Nat Genet 2021 Sep;53(9):1334-134756. GEO accession code: GSE176078
5. Breast cancer data from 17 individuals from the Array-Express database accession code: E-MTAB-8107. Described in (82) Qian, Junbin, *et al. “A pan-cancer blueprint of the heterogeneous tumor microenvironment revealed by single-cell profiling*.*”* Cell research 30.9 (2020): 745-76257.

## Acknowledgments

This work was supported by the Norwegian Research Council, Helse Sør-Øst, and University of Oslo through the Centre for Molecular Medicine Norway (187615, M.L.K and D.O), the Nor-wegian Cancer Society (214871, M.L.K.), the Norwegian Research Council (313932, M.L.K), and the Marie Skłodowska-Curie Postdoctoral Scientia Fellows program from the Faculty of Medicine at the University of Oslo, co-funded by the Euro-pean Union’s Horizon 2020 research and innovation programme under the Marie Skłodowska-Curie Actions Grant (801133, D.O and P.S).

## Supplementary Material

### Supplementary Figures

**Fig. S1.**
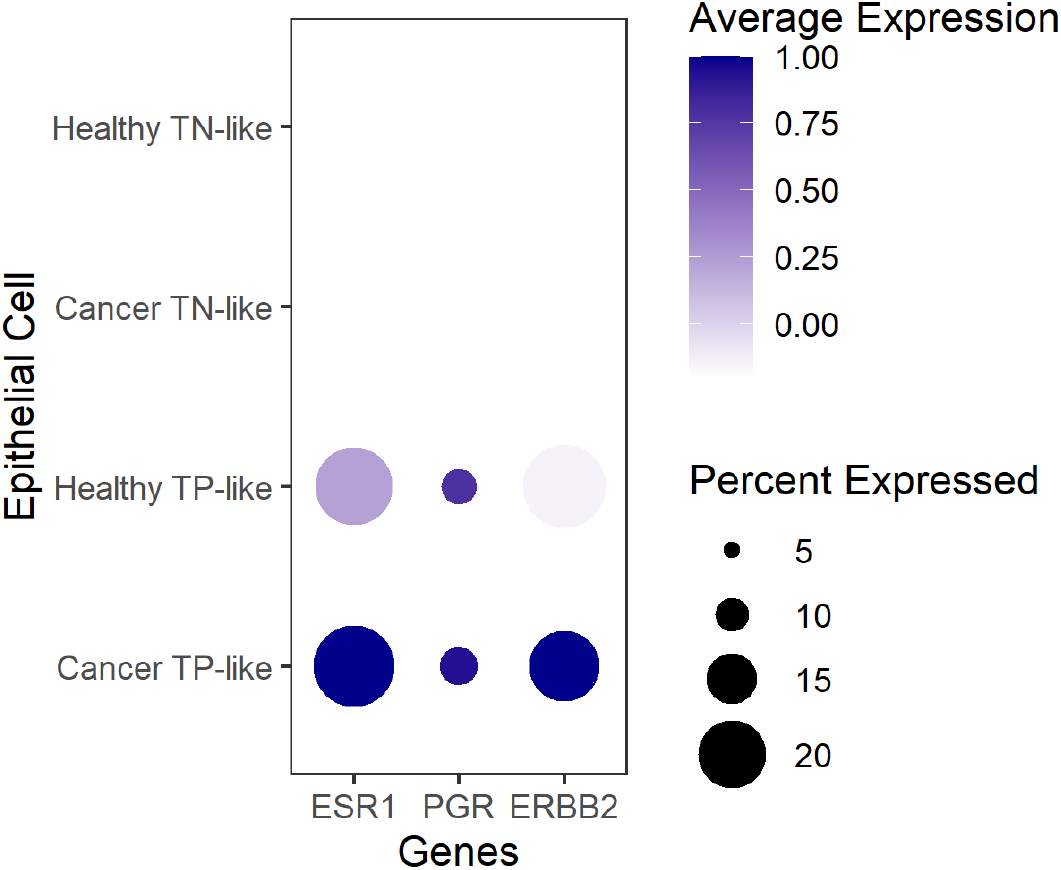
Comparisons of *ESR1, PGR*, and *ERBB2* expression levels across epithelial cells in cancer and healthy tissues. TP stands for Triple-Positive, and TN for Triple-Negative.

**Fig. S2.**
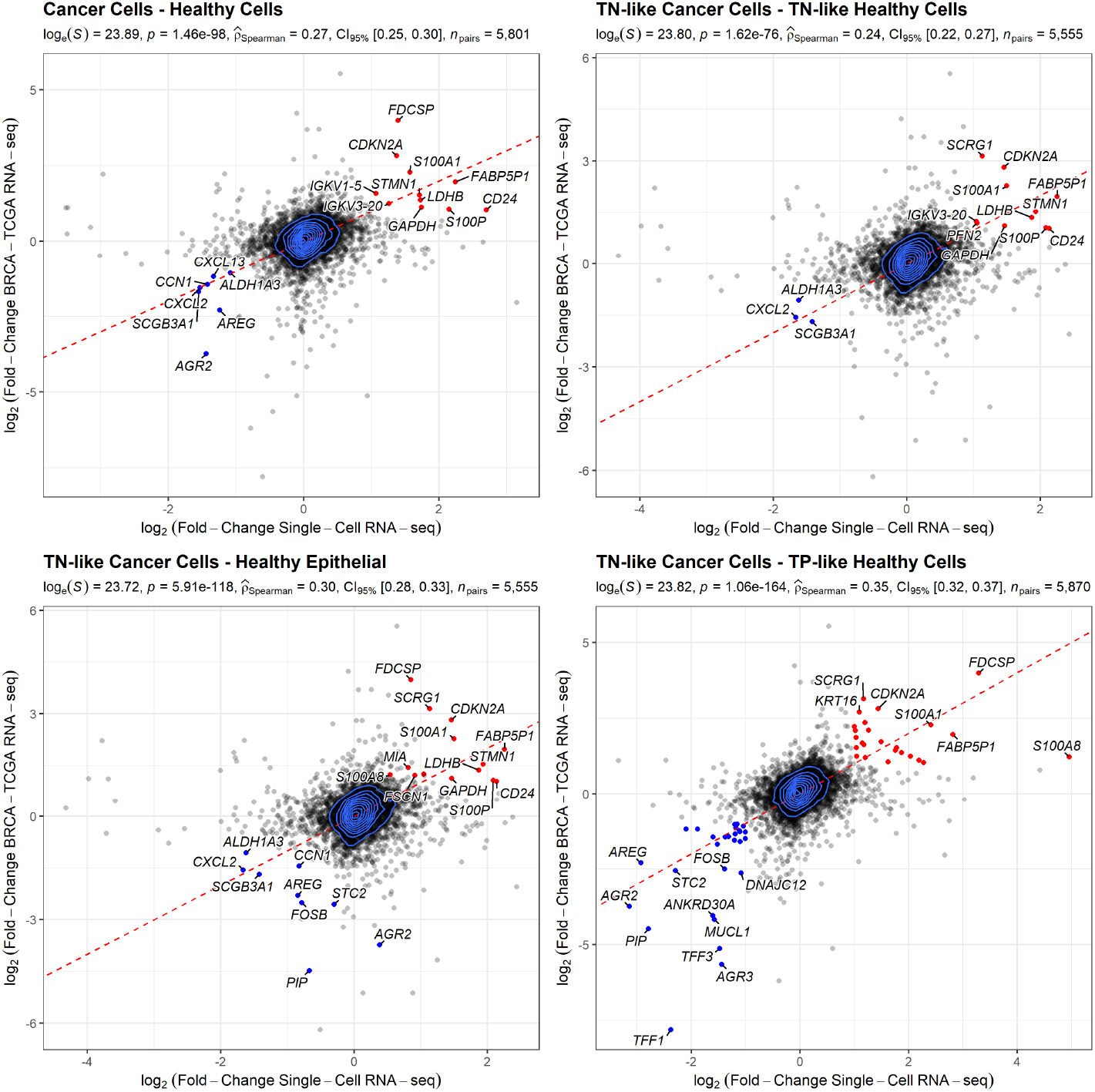
Comparisons of the transcriptional changes associated with TNBC at the single-cell and tissue level across different subpopulations of cells. Each dot represents a gene. Dots are color coded, in red if the log_2_ fold-change is larger than 1 and in blue if the log_2_ fold-change is smaller than −1. TP = Triple-Positive, TN = Triple-Negative.

**Fig. S3.**
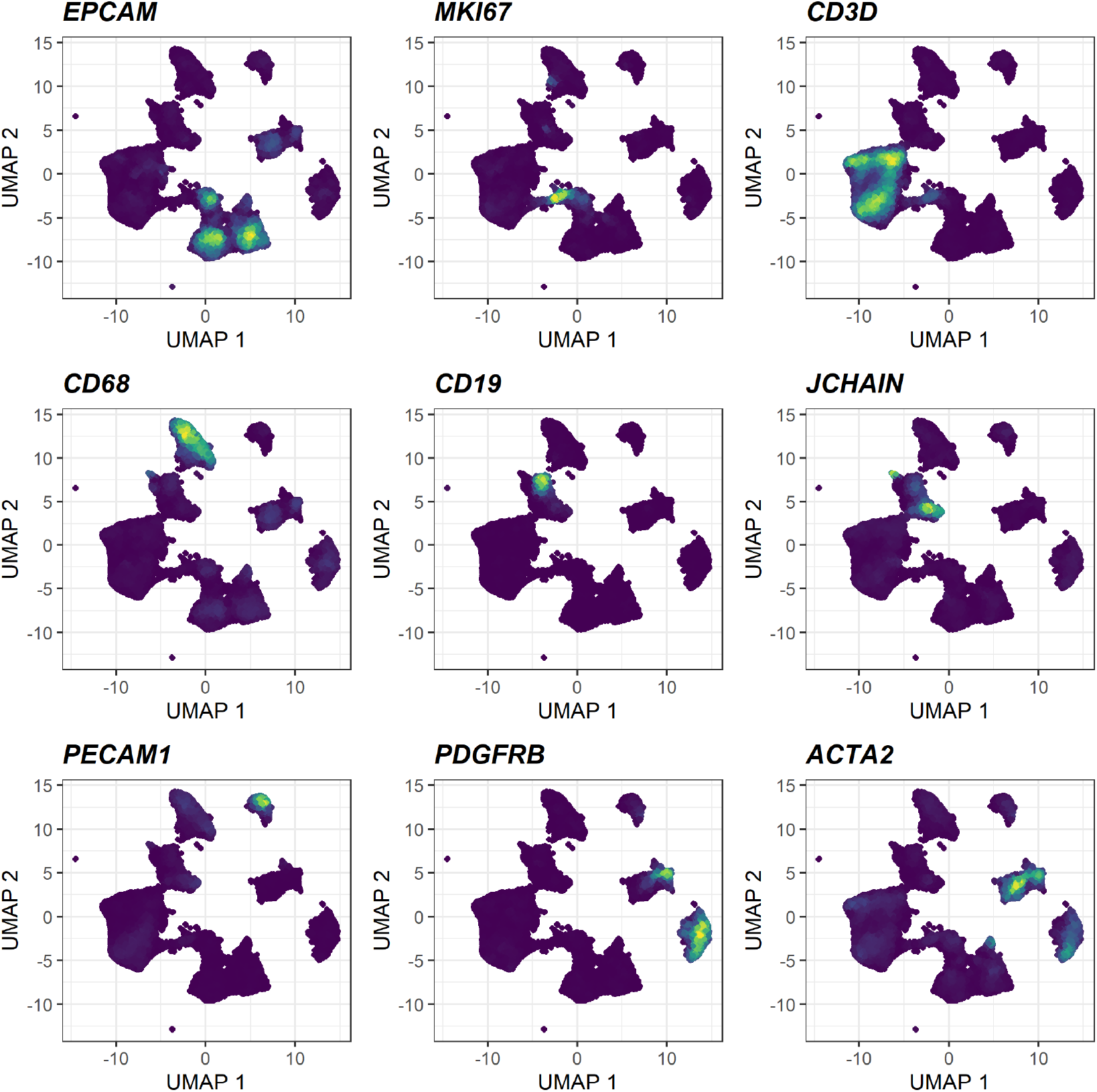
Expression of markers for epithelial cells (*EPCAM*), proliferating cells (*MKI67*), T cells (*CD3D*), myeloid cells (*CD68*), B cells (*MS4A1*), plasmablasts (*JCHAIN*), endothelial cells (*PECAM1*), mesenchymal cells (fibroblasts/perivascular-like cells; *PDGFRB*), and muscular cells (*ACTA2*).

**Fig. S4.**
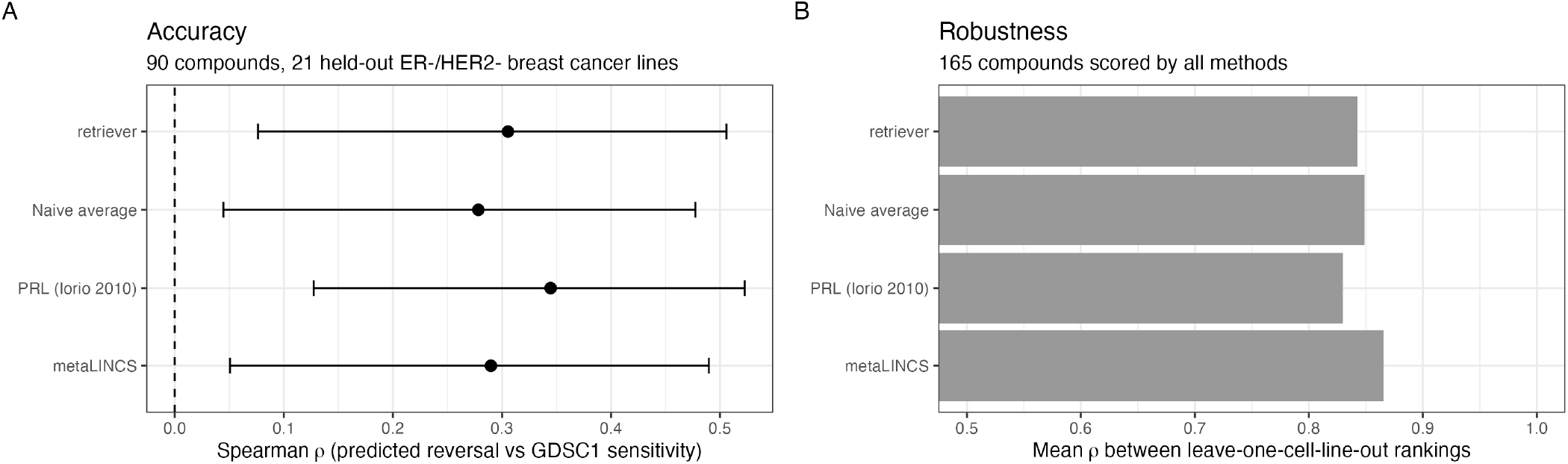
Benchmark of *retriever* against metaLINCS, the Prototype Ranked List (PRL), and naive averaging. **(A)** Spearman correlation between predicted reversal scores and GDSC1 drug sensitivity in 21 TNBC cell lines not profiled by LINCS-L1000 (90 compounds; 95% bootstrap confidence intervals). **(B)** Mean correlation between the rankings obtained from each pair of TNBC cell lines (165 compounds).

**Fig. S5.**
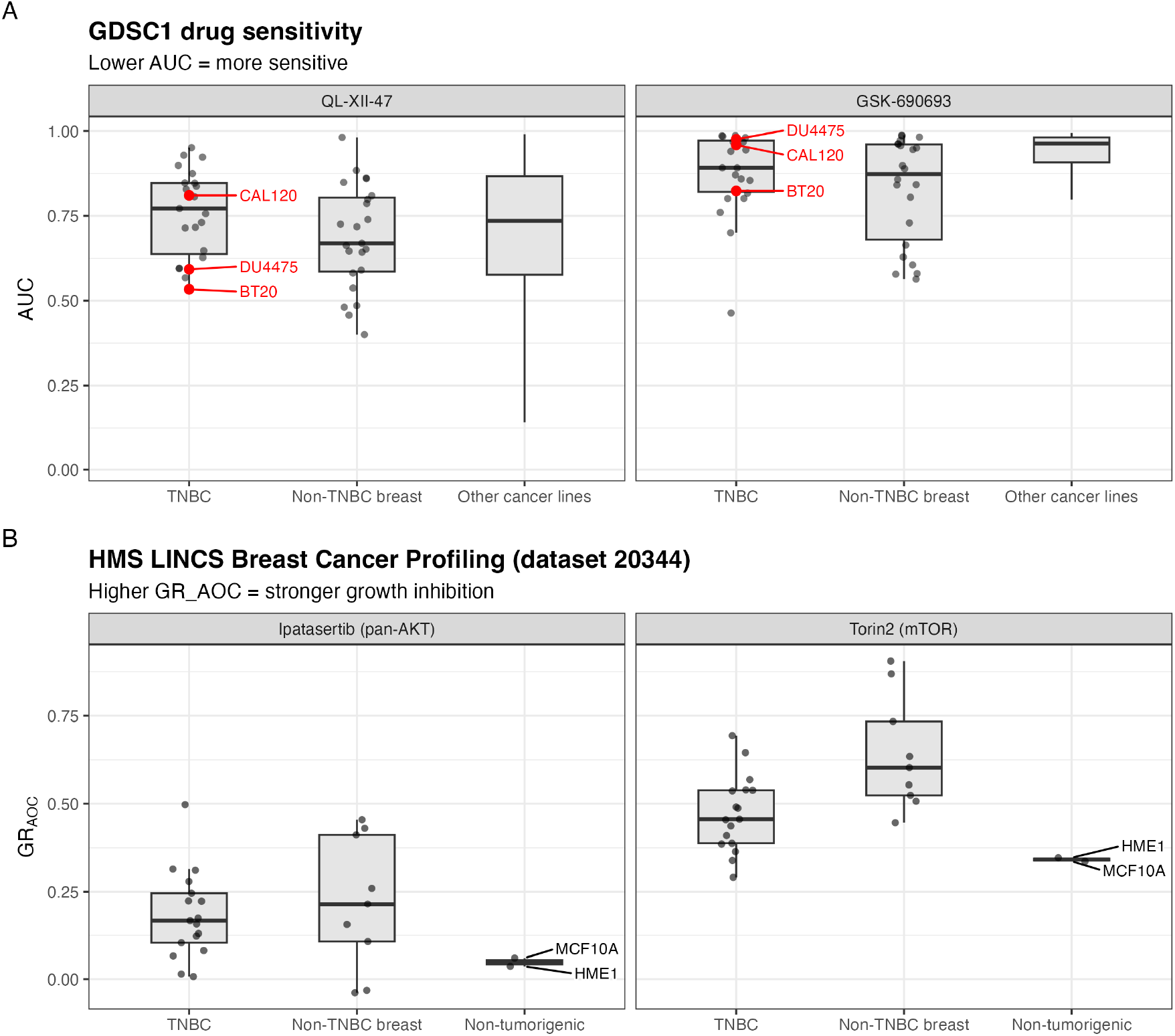
Public drug sensitivity data. **(A)** GDSC1 AUC for QL-XII-47 and GSK-690693 in TNBC and other breast cancer cell lines and in all other cancer cell lines; lower AUC indicates higher sensitivity. The cell lines tested in this study are highlighted in red. **(B)** Growth-rate inhibition (GR_AOC_) for ipatasertib and Torin2 in TNBC, non-TNBC, and non-tumorigenic breast cell lines (HMS LINCS dataset 20344); higher values indicate stronger inhibition.

**Fig. S6.**
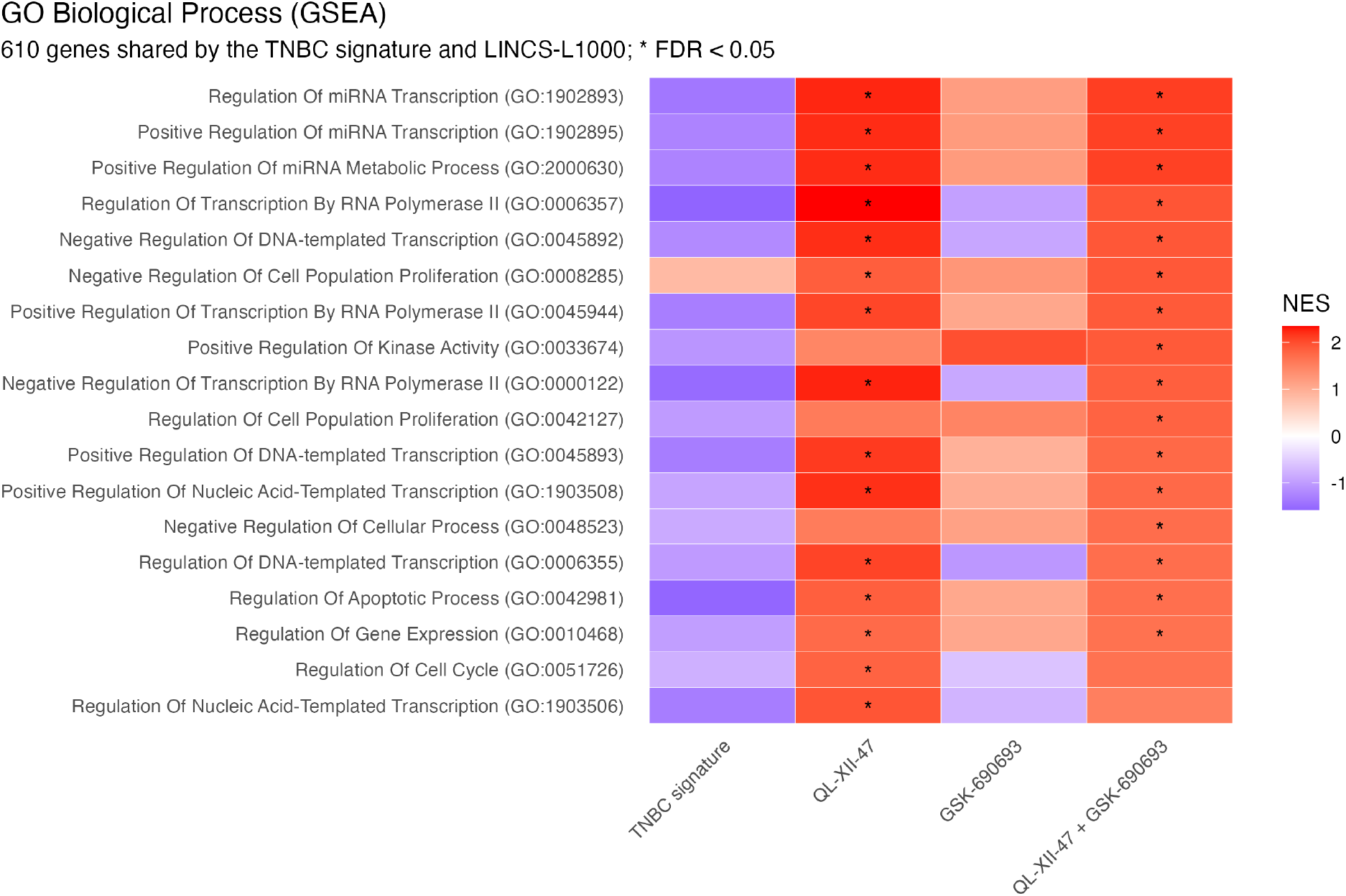
GO Biological Process enrichment (NES) of the TNBC signature and of the QL-XII-47, GSK-690693, and combination profiles on the 610 genes they share. Terms significant for at least one drug profile are shown; asterisks mark FDR *<* 0.05.

### Supplementary Tables

| Cancer Hallmark | FDR | Leading Genes |
|---|---|---|
| Oxidative Phosphorylation | 1.76 × 10<sup>−4</sup> | *LDHB, IDH2, SLC25A5, ATP5F1C, COX6C, PHB2, ATP6V0C,* and *UQCRH* |
| Interferon Alpha Response | 6.10 × 10<sup>−3</sup> | *CD74, IFITM1, IFI27,* and *LY6E* |
| Myc Targets | 4.24 × 10<sup>−2</sup> | *RPLP0, PABPC1, PHB2,* and *PPIA* |
| TNFα Signaling via NF-κB | 2.95 × 10<sup>−19</sup> | *EDN1, BTG1, GADD45B, TSC22D1, CEBPD, CXCL1, SOD2, AREG, CXCL2, NFKBIA, MARCKS, KLF6, MAFF, CCL2, FOSB, CCN1*, *IL6ST, IER5, ATF3,* and *IER3* |
| Estrogen Response | 6.54 × 10<sup>−8</sup> | *UGCG, KRT18, ELOVL5, STC2, HSPB8, SLC39A6, TFF3, PMAIP1, TFF1, IL6ST,* and *AREG* |
| UV Response Up | 6.54 × 10<sup>−8</sup> | *NFKBIA, BTG1, HNRNPU, HMOX1, FOSB, TMBIM6, IL6ST, SOD2, CXCL2,* and *ATF3* |
| Apoptosis | 6.54 × 10<sup>−8</sup> | *WEE1, KRT18, GADD45B, TNFRSF12A, PMAIP1, HMOX1, EMP1, SOD2, ATF3,* and *IER3* |
| Hypoxia | 3.45 × 10<sup>−7</sup> | *KLF6, BTG1, HSPA5, STC2, MAFF, HMOX1, ADM, CCN1, ATF3,* and *IER3* |
| Unfolded Protein Response | 9.36 × 10<sup>−5</sup> | *HSPA5, STC2, DNAJB9, CCL2, ATF3*, and *ATF4* |
| IL-6/JAK/STAT3 Signaling | 2.59 × 10<sup>−4</sup> | *TNFRSF12A, HMOX1, CXCL1, CXCL13,* and *IL6ST* |
| Inflammatory Response | 1.57 × 10<sup>−3</sup> | *NFKBIA, EDN1, KLF6, CXCL8, CCL2*, and *ADM* |
| Androgen Response | 4.62 × 10<sup>−3</sup> | *TSC22D1, ELOVL5, DNAJB9,* and *SLC38A2* |

**Table S1**. Hallmark MSigDB signatures associated with the differentially expressed genes observed in triple-negative breast cancer (TNBC). False Discovery Rate (FDR) computed using Gene Set Enrichment Analysis (GSEA).

**Table S2:** (CSV File) Computed robust disease-specific response profiles to 151 compounds across three triple-negative breast cancer cell lines (TNBC)

**Table S3:** (CSV File) Computed 11,325 compound combinations response profiles across three triple-negative breast cancer cell lines (TNBC)

**Table S4:** (CSV File) Ranked compounds based on their predicted potential to reverse the disease state (TNBC) towards a healthy state

**Table S5:** (CSV File) Ranked compound combinations based on their predicted potential to reverse the disease state (TNBC) towards a healthy state

**Table S6:** (CSV File) Cellular viability measurements among three distinct breast cancer cell lines following treatment with 0.6 *µ*M QL-XII-47, 0.8 *µ*M GSK-690693, and the combination of both compounds under same concentration

**Table S7:** (CSV File) Ranked compounds for each sample based on their predicted potential to reverse the disease state (TNBC) towards a healthy state

**Table S8:** (CSV File) Ranked compound combinations for each sample based on their predicted potential to reverse the disease state (TNBC) towards a healthy state

**Table S9:** (CSV File) Genes differentially expressed between TNBC-like and healthy triple-positive-like epithelial cells, with log_2_ fold-change, FDR, and direction

